# SNAPS: A toolkit for studying cell surface shedding of diverse transmembrane receptors

**DOI:** 10.1101/436592

**Authors:** Amanda N. Hayward, Eric J. Aird, Wendy R. Gordon

## Abstract

Proteolysis of transmembrane receptors is a critical cellular communication mechanism dysregulated in many diseases, yet decoding proteolytic regulation mechanisms of the estimated 400 receptors shed from the cell surface has been hindered by difficulties in controlling stimuli and unknown fates of cleavage products. Notch proteolytic regulation is a notable exception, where decades of study have revealed that intercellular forces drive exposure of a cryptic protease site within a juxtamembrane “proteolytic switch” domain to activate transcriptional programs inside the cell. Thus, we created a Synthetic Notch Assay for Proteolytic Switches (SNAPS) that exploits the modularity and unequivocal input/response of Notch proteolysis to screen surface receptors for other putative proteolytic switches. Here, we identify several new proteolytic switches among receptors with structural homology to Notch. We demonstrate that SNAPS can detect shedding in chimeras of diverse cell surface receptors, leading to new, testable hypotheses. Finally, we establish that the assay can be used to measure modulation of proteolysis by potential therapeutics.

## Introduction

Proteolysis of cell surface transmembrane proteins is a tightly regulated cellular mechanism that controls communication of cells with their extracellular environment. Diverse adhesion receptors, such as cadherins, as well as receptors that respond to soluble factors, such as receptor tyrosine kinases (RTKs), have been shown to be cleaved at sites close to the extracellular side of the membrane by metalloproteinases such as ‘A Disintegrin And Metalloproteinases’ (ADAMs) and matrix metalloproteinases (MMPs) (Miller, Sullivan, and Lauffenburger 2017; Kessenbrock, Plaks, and Werb 2010; McCawley and Matrisian 2001; Seals and Courtneidge 2003; White 2003), resulting in ectodomain shedding. In many of these receptors, ectodomain shedding is a prerequisite for further cleavage inside the membrane by the γ-secretase/presenilin protease complex in a process known as Regulated Intramembrane Proteolysis (RIP) (Brown et al. 2000; Selkoe and Wolfe 2007).

Proteolysis not only provides a mechanism to break cell-cell and cell-ECM contacts to modulate processes such as cell migration, but may also result in biologically-active fragments outside and inside of the cell, such as the intracellular fragment of Notch, which translocates to the nucleus and acts as a transcriptional co-activator (Bray 2006). Dysregulated proteolysis contributes to disease pathogenesis, for example, by causing accumulation of pathogenic fragments such as the amyloid beta peptide implicated in Alzheimer’s disease (Goate et al. 1991; Scheuner, Eckman, Jensen, Song, Citron, Suzuki, Bird, Hardy, Hutton, Kukull, Larson, et al. 1996), or removing epitopes required for normal cell communication (Groh et al. 2002; Kaiser et al. 2007; Boutet et al. 2009; Waldhauer et al. 2008). For instance, cancer cells evade the immune response by shedding MICA receptors (Groh et al. 2002; Kaiser et al. 2007; Boutet et al. 2009; Waldhauer et al. 2008), which are normally deployed to the cell surface in response to cellular damage.

Modulation of proteolysis is a heavily pursued therapeutic avenue, aiming to either inhibit proteases or prevent access to protease sites in a specific receptor. Many protease inhibitors have been developed but have failed clinically due to significant off-target effects (Dufour and Overall 2013; Vandenbroucke and Libert 2014; Turk 2006). Conversely, relatively few examples of modulating access to protease sites in specific receptors have been reported, despite the clinical success of the monoclonal antibody trastuzumab (Herceptin) that was found to act, in part, by blocking proteolysis of the receptor tyrosine kinase HER2 (Molina et al. 2001). Similarly, successful development of modulatory antibodies targeting proteolysis of Notch (Li et al. 2008; Aste-Amézaga et al. 2010; Wu et al. 2010; Tiyanont et al. 2013; Qiu et al. 2013; Agnusdei et al. 2014; Falk et al. 2012) and MICA (Ferrari de Andrade et al. 2018) receptors have recently been reported. However, though 8% of the annotated human transmembrane proteins are predicted to be shed from the surface (Tien, Chen, and Wu 2017), mechanisms of proteolytic regulation that inform development of specific modulators have remained elusive.

A relatively unique proteolytic regulation mechanism has recently come to light in which a stimulus alters protein conformation to induce exposure of a cryptic protease site. For example, the secreted von Willebrand factor (VWF) is cleaved in its A2 domain in response to shear stress in the bloodstream, which regulates blood clotting (Dong et al. 2002). Transmembrane Notch receptors also control exposure of a cryptic protease recognition site via the conformation of a juxtamembrane domain called the Negative Regulatory Region (NRR) (Gordon et al. 2015, 2007, 2009; Xu et al. 2015; Sanchez-Irizarry et al. 2004a)(Gordon et al. 2015, 2007, 2009; Xu et al. 2015) to trigger Notch signaling (Kopan and Ilagan 2009; Bray 2006) in response to ligand binding and subsequent endocytic forces (Parks et al. 2000; Gordon et al. 2015; Langridge and Struhl 2017). Crystal structures of the NRR (Gordon et al. 2007, 2009; Xu et al. 2015) reveal that the ADAM10/17 protease site is housed in a Sea urchin Enterokinase Agrin-like (SEA-like), with high structural homology to canonical SEA domains of mucins (Macao et al. 2006; Maeda et al. 2004) but lacking the characteristic autoproteolytic site. The NRR normally exists in a proteolytic cleavage-resistant state in which the protease site is buried by interdomain interactions between the SEA-like and its neighboring domain but can be switched to a protease-sensitive state when it undergoes a conformational change upon binding a ligand on a neighboring cell and subsequent endocytosis (Gordon et al. 2015; Parks et al. 2000; Sanchez-Irizarry et al. 2004b) or if it harbors disease-related mutations that destabilize the domain (Malecki et al. 2006; Gordon et al. 2009; Weng et al. 2004).

Notch’s proteolytic switch has been exploited to develop conformation-specific modulatory antibodies (Li et al. 2008; Aste-Amézaga et al. 2010; Wu et al. 2010; Tiyanont et al. 2013; Qiu et al. 2013; Agnusdei et al. 2014; Falk et al. 2012) and harnessed for synthetic biology applications (Morsut et al. 2016) to turn on transcription in response to any desired cell-cell contact. For example, Notch was engineered to respond to novel inputs and create custom responses (Roybal et al. 2016). This SynNotch system has been applied to CAR-T therapy to require dual antigen recognition for T-cell activation, increasing specificity. Thus, identification of additional proteolytic switches is of great interest. However, despite the knowledge that several cell-surface receptors harbor extracellular juxtamembrane domains with structural homology to Notch’s proteolytic switch (Pei and Grishin 2017) and that more than 100 receptors undergo a Notch-like proteolytic cascade (Brown et al. 2000; Selkoe and Wolfe 2007), other membrane resident proteolytic switches have not been identified, in large part due to difficulties in studying proteolysis in most receptors. For example, controlling the stimulus for receptors involved in homotypic interactions is difficult and the signaling pathways modulated by cleaved intracellular fragments may not be known.

A recent study showing that the known VWF proteolytic switch domain could functionally substitute for the Notch NRR to facilitate Notch signaling in certain contexts in Drosophila (Langridge and Struhl 2017) inspired us to ask if we could exploit Notch signaling to discover new proteolytic switches. We created a Synthetic Notch Assay for Proteolytic Switches (SNAPS) that harnesses the modularity and precise control of Notch signaling (Malecki et al. 2006; Gordon et al. 2015; Roybal et al. 2016) to screen protease site-containing juxtamembrane domains of diverse cell-surface receptors for their ability to functionally substitute for Notch’s proteolytic switch and induce transcription in response to cell-cell contact. SNAPS uses the native Notch ligand-binding interaction with DLL4 as the input and the Gal4 transcriptional response as the output (Fig. 1A). Here, we find that proteolysis regions of several receptors with structural homology to Notch can substitute for the Notch “proteolytic switch” and facilitate signaling in response to cell contact. Moreover, the assay can be used to detect shedding of diverse receptors such as RTKs, CD44, and cadherins. Finally, we demonstrate that the assay can be used to screen modulators of proteolysis.

**Fig. 1:**
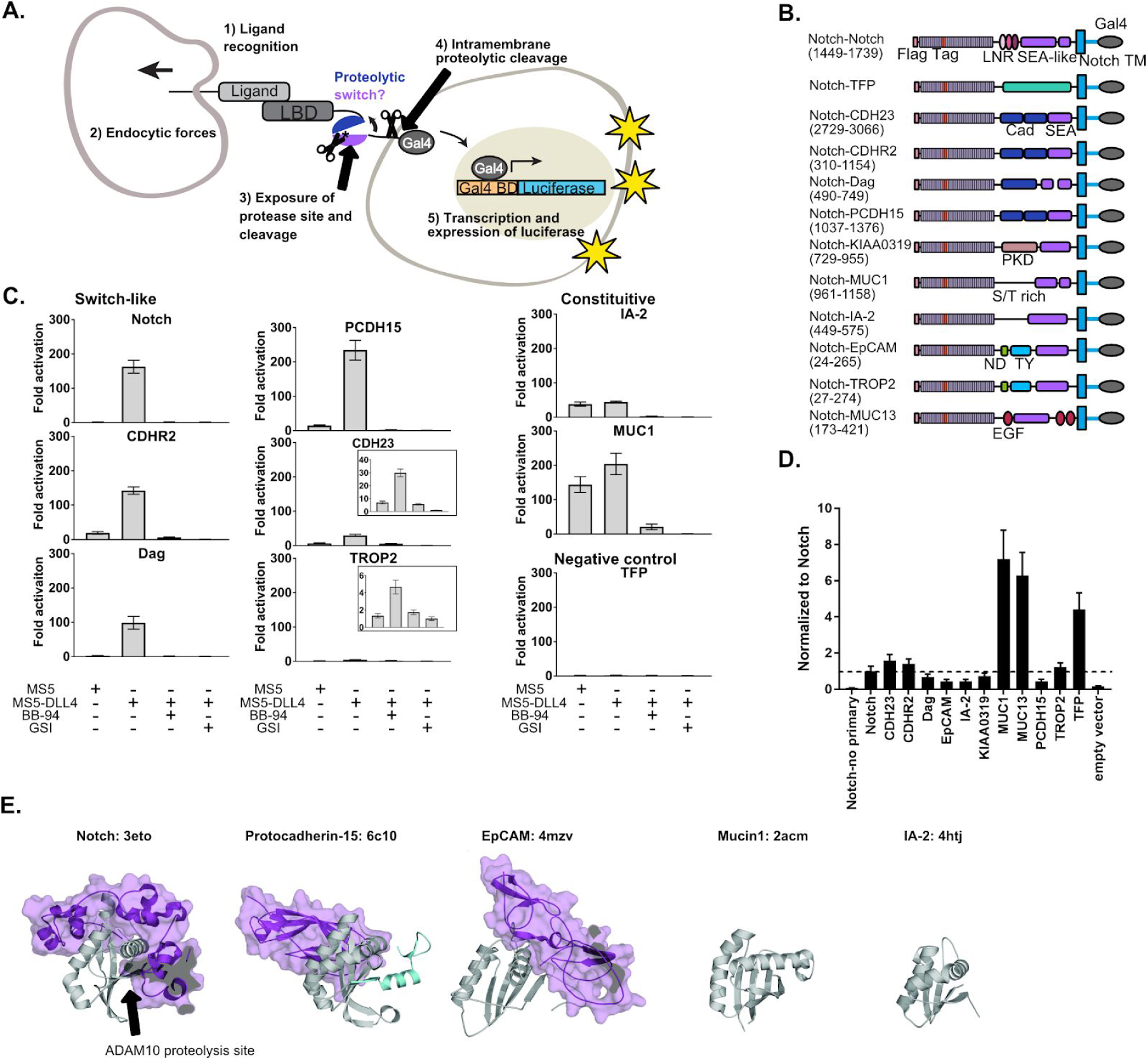
SEA-like domains cooperate with adjacent domains to behave as proteolytic switches. (**A**) Schematic of Synthetic Notch Assay for Proteolytic Switches (SNAPS). Cells co-expressing Flag-Notch-X-Gal4 chimeras, where X is a putative proteolysis region of another receptor, and luciferase reporter constructs are co-cultured with DLL4 ligand-expressing cells to induce Notch activation and expression of luciferase. (**B**) Schematic of chimeric constructs utilized in the signaling assay. Protein domains are color coded and labeled below. Amino acid ranges used for each construct are in parentheses under the names. Note that Notch’s SEA-like domain is also referred to as the Heterodimerization Domain (HD) in the literature. Abbreviations used: Cad: cadherin. EGF: Epidermal growth factor. LBD: Ligand binding domain. LNR: Lin-12 Notch-like repeats. ND: N-terminal domain. PKD: polycystic kidney disease domain. S/T rich: serine-threonine rich. TFP: Teal fluorescent protein. TM: transmembrane domain. TY: thyroglobulin type-1A domain. (**C**) Luciferase reporter gene activity profile of Notch and Notch chimera constructs co-cultured with MS5 cells or MS5 cells stably expressing DLL4. BB-94=Batimastat (pan-metalloproteinase inhibitor) GSI= Compound E (γ-secretase inhibitor). Error bars represent the SEM of triplicate measurements. (**D**) Cell surface ELISA of Notch and Notch chimera constructs. Anti-Flag primary and goat anti-mouse HRP secondary antibodies were used to detect cell surface expression levels of each chimera. The horizontal dotted line corresponds to Notch expression levels. Error bars represent the SEM of triplicate measurements. (**E**) Structures and PDB IDs of SEA-like domains (grey) with applicable adjacent domains (purple). SEA-like domains were structurally aligned to the Notch SEA-like domain.

## Results

### SEA-like domains cooperate with adjacent domains to behave as proteolytic switches

To determine if receptors bearing juxtamembrane domains predicted to be structurally homologous to Notch could also function as proteolytic switches, we created chimeric receptors in which we replaced the Notch NRR proteolytic switch domain with SEA and SEA-like domains from other cell surface receptors and included any previously characterized tandem N-terminal domains (Fig. 1B, Supplementary Table 1). We also made a negative control chimera where the NRR was replaced by the fluorescent protein mTFP. We hypothesized that other putative proteolytic switches could functionally substitute for the Notch NRR and initiate a transcriptional response in response to contact with a cell expressing DLL4. Then, chimeric constructs together with Gal4-responsive and control luciferase reporter constructs were transfected into U2OS cells, co-cultured with cells stably expressing Notch ligands, and luciferase activity measured in a high-throughput format.

Surprisingly, we found that the ECM receptor dystroglycan and two protocadherins involved in intercellular adhesion, Protocadherin-15 (PCDH15) and Cadherin-related protein 2 (CDHR2), could functionally substitute for Notch’s NRR (Fig. 1C). These chimeric receptors signaled robustly only in the presence of cells expressing DLL4, and the signal was abrogated by both a global metalloproteinase inhibitor (BB-94) and an inhibitor of the subsequent intramembrane γ-secretase cleavage event (γ-secretase inhibitor; GSI). The putative cell adhesion molecules Trop2 and Cadherin-23 (CADH23) displayed a more moderate signaling activity in response to DLL4. Interestingly, these receptors all contain a SEA or a SEA-like domains in tandem with an N-terminal domain. On the other hand, SEA/SEA-like domains without a structured neighboring domain, such as Mucin-1 (MUC1) and receptor tyrosine phosphatase-related islet antigen 2 (IA-2), exhibited a high level of proteolysis even in the absence of DLL4, suggesting they contain a constitutively exposed protease site in the context of this assay.

A few chimeras showed very little signal in the assay (Fig. S1), suggesting a lack of proteolysis or lack of cell-surface expression. To further probe the receptors exhibiting low levels of activation in the signaling assay, we performed a cell-surface ELISA assay. Briefly, Flag-tagged Notch chimera constructs were transfected into U2OS cells, fixed, stained, and cell-surface levels quantified by measuring HRP activity. Most of the chimeras lacking signaling activity also expressed at lower levels than Notch, suggesting defects in expression or trafficking due to incorrect choice of domain termini. Our negative control mTFP chimera and MUC13 expressed substantially better than Notch (Fig. 1D), suggesting lack of response in the signaling assay is due to an absence of proteolysis in the assay. In contrast, the ELISA showed that IA-2 expressed at much lower levels than Notch yet exhibited robust constitutive signaling activity. We reasoned that high rates of shedding could result in apparently low cell-surface levels in the ELISA assay, so we repeated the ELISA assay with the addition of the metalloproteinase inhibitor BB-94. Indeed, IA-2 surface expression was substantially increased in the presence of BB-94 (Fig. S2A), while surface levels of other receptors that exhibited constitutive signaling activity were not drastically affected by inhibitor treatment. This suggests that IA-2 undergoes much higher rates of proteolysis than the other proteins studied. Since we observed variable cell-surface levels of the receptors in the ELISA assay, we also performed titrations of the chimeric receptors in the SNAPS signaling assay to ensure that high surface level expression was not masking proteolytic switch like behavior (Fig. S3A and C). Most receptors showed decreasing signaling activity with decreasing concentration of receptor, as expected. Interestingly, IA-2 signal increased as receptor concentration decreased, perhaps related to its high expression levels and turnover rates.

Comparing solved structures of several SEA/SEA-like domain containing proteins reveals additional insights (Fig. 1E). SEA/SEA-like domains are colored grey with adjacent N-terminal domains in purple. In contrast to Notch, Protocadherin-15, and EpCAM, MUC1, and IA-2 do not have structured domains N-terminal to their SEA/SEA-like domain. This likely leads to enhanced conformational dynamics, resulting in an increase in protease site exposure and signaling. Though Notch and Protocadherin-15 exhibit similar conformational switch behavior in the signaling assay, Protocadherin-15’s N-terminal cadherin-like domain binds on the opposite face of the SEA-like domain than Notch’s neighboring Lin12 Notch Repeat domain, suggesting potentially different conformational switching propensities.

### SNAPS to probe MMP proteolysis of dystroglycan

Next, we aimed to validate the putative proteolytic switch behavior of a receptor / Notch chimera revealed by SNAPS (Fig. 2). Dystroglycan provides a critical mechanical link between the ECM and the actin cytoskeleton to help muscle cells withstand contraction and neural cells maintain the blood brain barrier (Barresi and Campbell 2006; Agrawal et al. 2006). It is post-translationally processed into two subunits, termed α‐ and β-dystroglycan in its SEA-like domain, akin to mucins.

**Fig. 2:**
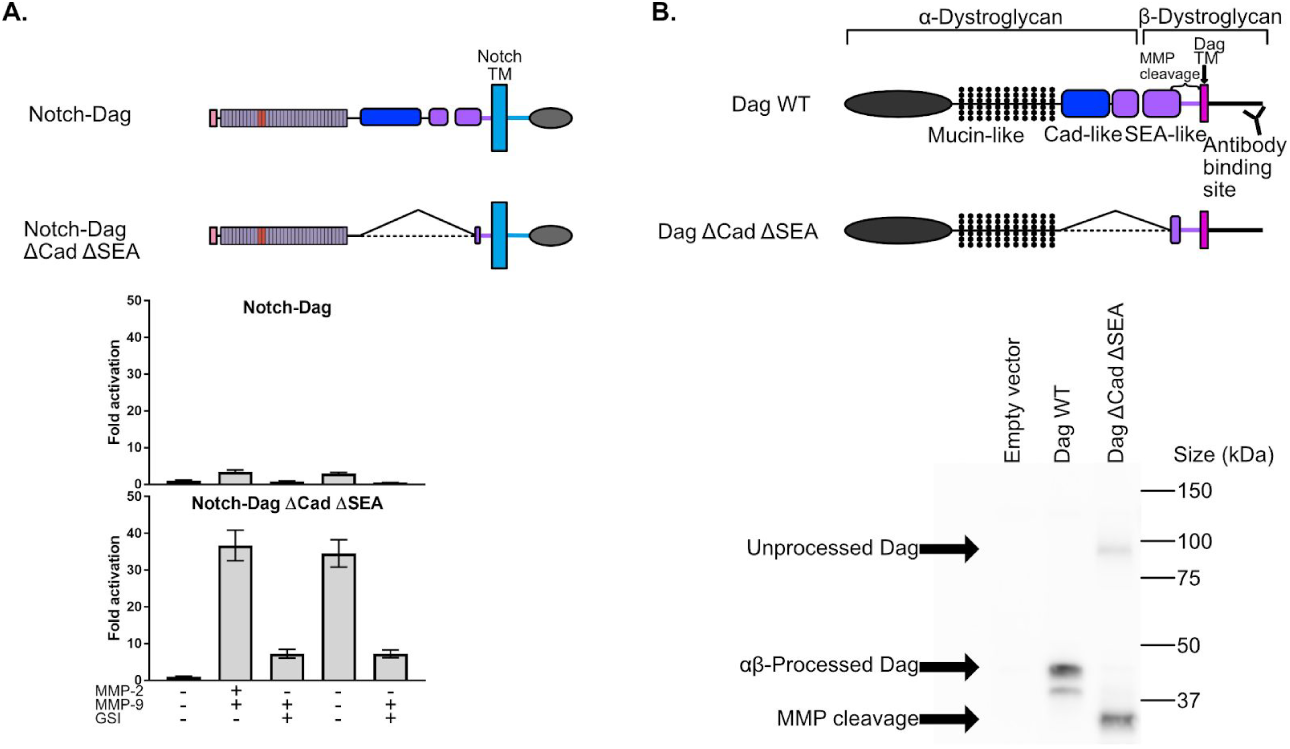
SNAPS to probe MMP proteolysis of dystroglycan. (**A**) Luciferase reporter gene activity of Notch-Dag chimeras containing intact proteolytic switch and truncated switch with constitutive MMP sites (ΔCadΔSEA) upon addition of MMPs. Error bars represent the SEM of triplicate measurements, normalization to no added MMP condition. (**B**) β-dystroglycan western blot of Cos7 cell lysates after transfection with empty vector, wild-type dystroglycan or dag ΔCadΔSEA. Bands for unprocessed, αβ-processed Dag, and MMP cleavage are denoted.

Cleavage of the 42 kDa β-dystroglycan by matrix metalloproteinases (MMPs) to a 31 kDa product is greatly enhanced in pathogenic states, such as muscular dystrophy and cancer (Agrawal et al. 2006; Matsumura et al. 2005; Singh et al. 2004). Thus, we asked whether receptors containing the entire proteolytic switch domain were more resistant to MMPs than a truncated proteolytic switch domain construct (ΔCadΔSEA) containing only the protease sites, in which proteolysis should be constitutive. We first used SNAPS to measure proteolysis induced by adding exogenous MMPs to Dystroglycan/Notch chimeras containing full-length and ΔCadΔSEA proteolytic switch domains. The chimeras containing intact proteolysis domains were indeed more resistant to MMPs than the chimera with constitutively exposed sites (Fig. 2A), exhibiting a modest 2-3 fold increase in basal proteolysis compared to a 30-fold increase in the ΔCadΔSEA Dag Notch chimera. We also saw a similar effect when adding MMP buffer containing APMA, a compound that activates MMPs (Fig. S4)

We next asked whether the truncated and intact proteolytic switch domains exhibited different sensitivities to proteolysis in the context of the non-chimeric, native dystroglycan receptor. Constructs were transfected into Cos7 cells, in which dystroglycan proteolysis has been previously studied, (Herzog et al. 2004)) and a Western blot of cell lysates was performed using a β-dystroglycan antibody to measure cleavage of the 42 kDa fragment to a 31 kDa fragment. As expected, wild-type dystroglycan with intact proteolytic switch shows low levels of the 31-kDa cleavage product compared to the substantial proteolysis observed for the mutant with constitutively exposed cleavage sites (Fig. 2B). These results suggest that the proteolytic switch-like behavior may be relevant to regulation of dystroglycan’s cleavage by MMPs in its native context, and that this assay can be used to further test hypotheses about regulation and potential modulation of proteolysis in dystroglycan.

### Shedding of diverse receptors detected by SNAPS

We next wanted to determine whether SNAPS could be used to detect membrane shedding of receptors that do not contain SEA-like domains. Proteolysis plays a major role in the function of cell surface receptors such as E-cadherin and RTKs (Merilahti et al. 2017; Okamoto et al. 1999; Katayama et al. 1994), and dysregulation of proteolysis in these receptors is linked to cancer pathogenesis (S. M. Brouxhon et al. 2014; Katayama et al. 1994; Okamoto et al. 1999; Arribas et al. 2011) and resistance to kinase inhibitor treatment (Miller et al. 2016; Colomer et al. 2000; Leitzel et al. 1995), for instance. Unlike the aforementioned receptors with putative protease sites housed in structured SEA/SEA-like domains, the protease sites responsible for receptor shedding in cadherins and RTKs map to a putatively unstructured region between a structured repeat and the transmembrane region (Franklin et al. 2004; D’Huyvetter et al. 2017; Cho et al. 2003; Harrison et al. 2011). These receptors (Fig. 3A) might be expected to have higher basal levels of proteolysis and proteolytic regulation mechanisms distinct from SEA-like domain containing receptors(Fig. 3B), however, an assay that can detect proteolysis in such receptors could provide a starting point to test hypotheses about other potential mechanisms to regulate shedding, such as disruption of dimerization interfaces. We first tested an E-cadherin chimera comprised of the two cadherin repeats closest to the transmembrane (Fig. 3B) either including or lacking the sequence containing putative cleavage sites between the terminal cadherin repeat and the membrane. The observed 10-fold increase in basal signaling (co-culture with MS5 cells) over Notch was abrogated by treatment with protease inhibitors and when the linker containing putative ADAM10 sites was removed, suggesting that the signal is due to shedding. SNAPS was able to measure basal levels of proteolysis that were generally higher (5 to 25 fold) than the SEA-like containing chimeras in most of the receptors we tested, including HER and TAM family RTKs, E-cadherin, MICA, and CD44 (Fig. 3B and Fig. S5A). Interestingly, several receptors exhibited significant increases in signaling when induced by Notch ligands, resembling proteolytic switches. These results suggest that exposure of cryptic protease sites might contribute to proteolytic regulation in these receptors (Fig. 3B), perhaps by altering conformations of dimers. Neither wild-type or mutated von Willebrand factor (VWF) A2 domain chimeras, which have been previously tested as functional replacements of the Notch NRR in *Drosophila* (Langridge and Struhl 2017), signaled in the context in this assay (Fig. S5A). We again performed receptor titrations and cell-surface ELISAs (Fig. S3B and 3D) and were encouraged that the receptors lacking structural homology to Notch also readily trafficked to the cell surface (Fig. S5B), demonstrating that SNAPS can be used to investigate shedding in a broad landscape of cell surface receptors.

**Fig. 3:**
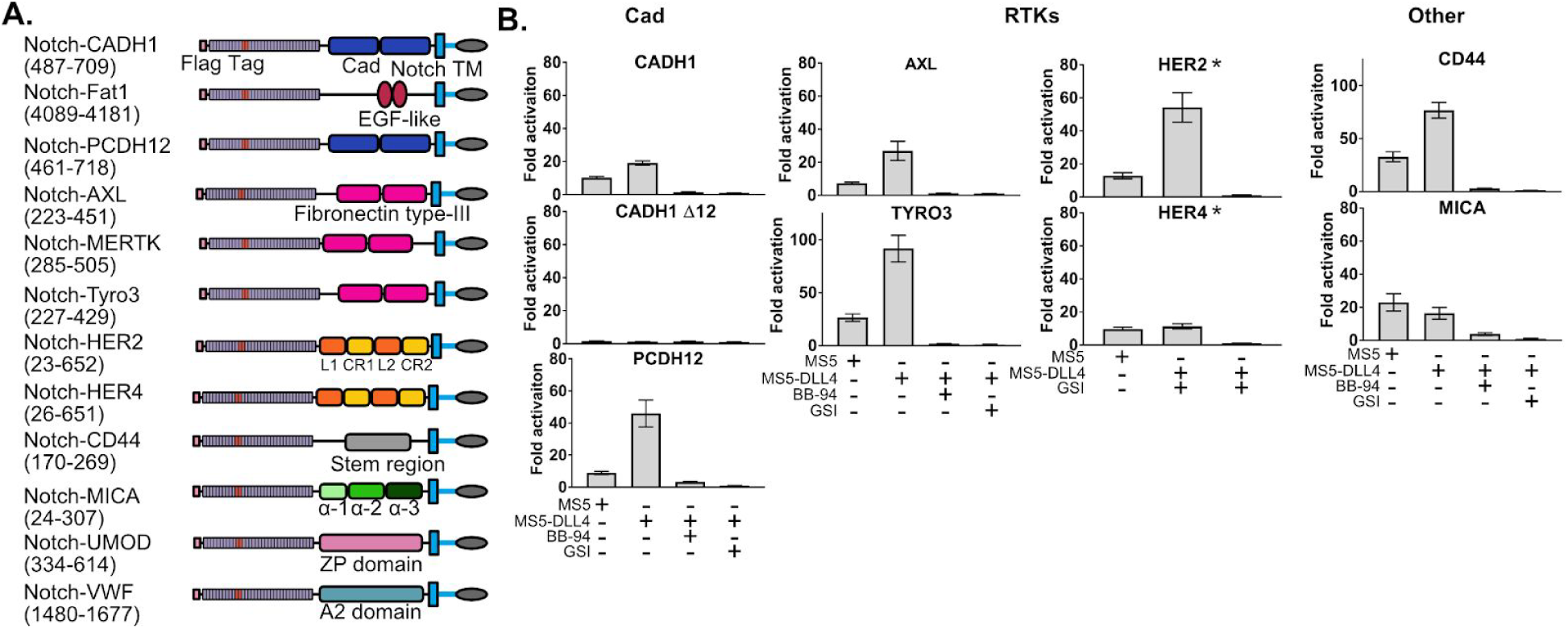
Shedding of diverse receptors detected by SNAPS. (**A**) Chimera constructs for proteins without SEA domains. Protein domains are color coded and labeled. (**B**) Luciferase reporter gene activity profile of Notch chimera constructs co-cultured with MS5 cells or MS5 cells stably transfected with DLL4, including treatment with BB-94 metalloprotease and GSI gamma secretase inhibitors as noted. Asterisked graphs denote experiments performed on different days and with 2ng DNA/well instead of 1ng/well of DNA. Error bars represent the SEM of triplicate measurements. CD44 and TYRO3 are shown with different scalebars due to high signal. Abbreviations:CADH1: E-cadherin. PCDH12: Protocadherin-12. CR: Cysteine rich. EGF: Epidermal growth factor. AXL: Tyrosine-protein kinase receptor UFO. HER2: human epidermal growth factor receptor 2.HER4: Human epidermal growth factor receptor 4. MICA: MHC class I polypeptide related sequence A. Tyro3: Tyrosine-protein kinase receptor TYRO3. Cad: Cadherin. UMOD: Uromodulin. VWF: Von Willebrand Factor. L: Leucine-rich. ZP: zona pellucida.

### SNAPS can be used to screen for proteolysis modulators

We next reasoned that SNAPS could provide a powerful means to screen for receptor-specific modulators of proteolysis. For example, the Notch signaling assay was used to screen activating and inhibitory therapeutic antibodies targeting the Notch receptor (Li et al. 2008; Aste-Amézaga et al. 2010; Wu et al. 2010). The monoclonal antibody trastuzumab (Herceptin), used to treat HER2+ breast cancer (Pegram et al. 1998; Baselga et al. 1996), has been shown to block proteolysis of the HER2 receptor tyrosine kinase as part of its mechanism of action (Molina et al. 2001). Therefore, we tested whether Herceptin could modulate the basal proteolysis of HER2 observed in the Notch-HER2 chimeras, in which the Notch NRR is replaced with the ectodomain of HER2.

We treated HER2-chimera expressing cells with increasing concentrations of Herceptin or an IgG control. We observed reproducible and statistically significant decreases in proteolysis in cells treated with Herceptin as compared to the IgG control (Fig. 4A). Proteolysis was reduced up to 40%. This robust effect demonstrates the potential utility of the chimeric assay in drug screening.

**Fig. 4:**
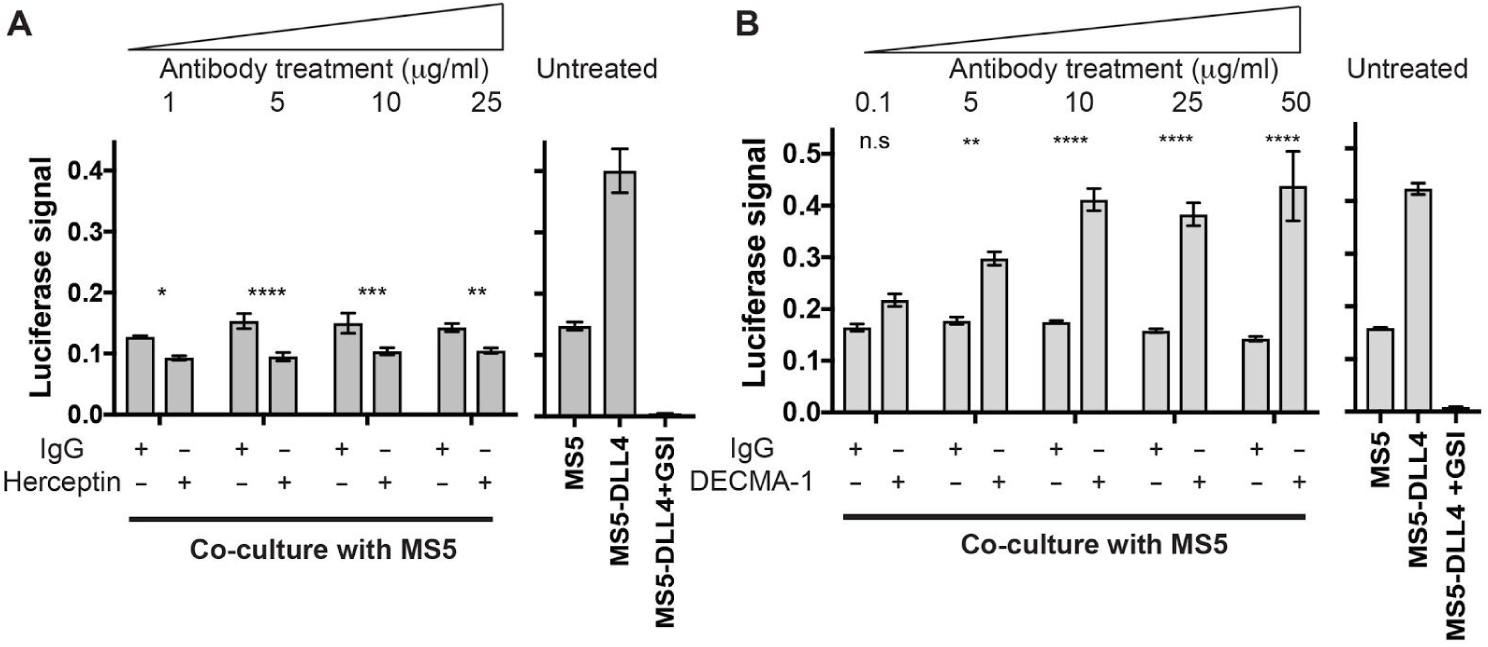
SNAPS can be used to screen for proteolysis modulators. (**A**) Left panel shows effects of Herceptin on basal signaling of HER2-Notch chimeras (i.e. co-culture with MS5 cells). HER2-Notch chimera expressing cells co-cultured with MS5 cells were treated with 1-25 ug/ml Herceptin or IgG control. Right panel shows untreated cells in co-culture with MS5 or MS5-DLL4 cells +/− GSI for reference. Error bars represent the SEM of triplicate measurements (**B**) Left panel shows effects of DECMA-1 on basal signaling of Cad4-5-Notch chimeras. Cad4-5-Notch chimeras co-cultured with MS5 cells were treated with 0.1-50 ug/ml of DECMA-1 or IgG control. Right panel shows untreated chimera basal and ligand-induced proteolysis shown for reference. Error bars represent the SEM of triplicate measurements. Statistical significance was determined with a two-way ANOVA followed by a post hoc Bonferroni test. ****: p<0.0001, ***: p<0.001, **:p<0.01 *: p<0.02

We next tested the effects of DECMA-1 on proteolysis of the Cadherin-Notch chimera. DECMA-1 is a function-blocking E-cadherin antibody known to break cell-cell contacts and reduce tumorigenesis in mice (Sabine M. Brouxhon et al. 2013). However, its mechanism of breaking cell-adhesions has remained elusive; the antibody binds to E-cadherin at the interface of the last two cadherin repeats (EC4 and EC5) near the membrane, but the N-terminal repeats EC1 and EC2 are responsible for the homotypic interactions presumed to be disrupted by the antibody. We hypothesized that DECMA-1 might affect E-cadherin shedding since the antibody epitope maps to the “proteolysis region” of E-cadherin.

In the absence of antibody or when treated with IgG control antibodies, the Notch-cadherin chimera, in which cadherin repeats EC4 and EC5 have replaced the Notch proteolytic switch, displays a moderate level of constitutive proteolysis and a 2-fold increase in activity in response to DLL4 expressing cells (Fig. 4B). When the cells are treated with DECMA-1, we observe a dose-dependent increase in the basal level of signaling, in comparison to control antibody, almost to the level of ligand induced signaling. The apparent EC50 of DECMA-1 measured by the assay is ∼0.8 ug/mL (Fig. S6). These data suggest that the mechanism of DECMA-1 breaking adhesive contacts could, in part, be due to increased shedding of the receptor from the membrane.

## Discussion

Notch’s proteolytic switch has been exploited to develop conformation-specific modulatory antibodies and harnessed for synthetic biology applications to turn on transcription in response to any desired cell to cell contact (Li et al. 2008; Aste-Amézaga et al. 2010; Wu et al. 2010; Tiyanont et al. 2013; Qiu et al. 2013; Agnusdei et al. 2014; Falk et al. 2012; Roybal et al. 2016; Gordon et al. 2015). Thus we created SNAPS utilizing well-understood stimuli and responses of Notch signaling to identify novel proteolytic switches and probe shedding of a wide range of cell-surface receptors. Using this assay, we find that Notch’s mechanism of proteolytic regulation via conformational control of a cryptic protease site is not a unique phenomenon and is rather a potentially common mechanism of control for several SEA-like domain-containing receptors that share structural homology to Notch. Moreover, shedding of transmembrane proteins such as HER2, AXL, CD44, and PCDH12 was detected with the assay, enabling new hypothesis generation about proteolytic regulation and modulation. SNAPS can also be used to screen for modulators of proteolysis; we observe that Herceptin treatment causes significant decreases in basal proteolysis of HER2, while DECMA-1 treatment results in substantial increases in basal proteolysis of E-cadherin. These results reveal new mechanistic insights into DECMA-1’s function in breaking cellular adhesions.

### New proteolytic switches for synthetic biology

Our studies revealed that most receptors containing juxtamembrane SEA-like domains are robustly shed from the cell-surface and that several of them behave as proteolytic switches, only becoming sensitive to proteolysis when “induced” with forces derived from cell-cell contact. We were struck by the fact that almost all of the receptors harboring SEA / SEA-like domains in tandem with neighboring domains showed a similar switch-like behavior in the intercellular signaling assay, despite having neighboring domains with very different predicted structural characteristics. In Notch, the neighboring LNR domain is disulfide-rich and binds calcium, with little to no secondary structure (Fig. 1E). Dystroglycan and the Protocadherins have neighboring cadherin-like domains, characterized by high β-strand content, while EpCAM and Trop2 have a cysteine-rich thyroglobulin domain. The existing crystal structures of several of these domains also reveal differential modes of interaction with the SEA/SEA-like domain. For example, in the EpCAM and NRR structures, the neighboring domain contacts the α-helix in close proximity to the β-strand containing putative proteolytic sites. In contrast, the cadherin-like domain interacts with the opposite face of the SEA-like domain in the Protocadherin-15 structure (Fig. 1E). These different modes of interdomain interactions suggest that the proteolytic switches may have different propensities to “switch on” as well as potentially different requirements for direction of applied force. Future studies probing comparative anatomy of putative proteolytic switches may reveal whether the structural differences are a consequence of cellular context; e.g. receptor involved in intercellular versus ECM interactions. On the other hand, the SEA-like domains lacking structured neighboring domains exhibit constitutive signaling, likely due to a more dynamic domain where protease site exposure occurs more frequently.

Synthetic biology applications that aim to induce transcription of a desired gene in response to cell to cell contact might benefit from proteolytic switches with different characteristics from the NRR of Notch. For example, in CAR-T applications, perhaps a switch that requires more “force” to switch on could be used to distinguish an epitope that is presented on a tumor with a stiff ECM compared to a normal cell. Moreover, the smaller and structurally simpler design of the Cadherin-like neighboring domains of dystroglycan and protocadherin-15 might permit more facile trafficking and expression for certain applications. Finally, constitutively proteolyzed MUC1 and IA2 exhibit much higher expression/rates of proteolysis and may provide opportunities for engineering more robust switches when paired with a neighboring domain.

### Targeting proteolytic switches and shedding with therapeutics

Notch’s proteolytic switch has been specifically targeted with both inhibitory and activating antibodies, suggesting that similar strategies could be successful for other receptors harboring proteolytic switches that are dysregulated in disease. While the proteolytic switches identified here need to be validated to determine if exposure of cryptic protease sites is physiologically relevant, we provided preliminary validation that dystroglycan protease sites may be conformationally controlled in the native receptor. Moreover, high levels of MMPs and thus dystroglycan cleavage have been observed in muscle biopsies of muscular dystrophy patients (Matsumura et al. 2005), and treatment of muscular dystrophy mouse models with broad spectrum metalloprotease inhibitors has been shown to ameliorate symptoms in a muscular dystrophy mouse model (Kumar et al. 2010). The dystroglycan proteolytic switch might offer a receptor-specific therapeutic target in diseases where MMP cleavage is dysregulated. Moreover, SNAPS was also able to detect shedding and modulation of shedding in receptors that do not contain SEA-like domains, suggesting that the assay can provide a platform to screen modulators of shedding of diverse receptors.

### Exposure of cryptic protease site may be a common mechanotransduction mechanism

In this study, mechanical force derived from intercellular contact is applied to cell-surface receptors to identify cryptic protease sites. While mechanical force may not play a role in the function of some receptors studied here, several of the receptors probed have been implicated in mechanosensing. Like Notch (Gordon et al. 2015; Parks et al. 2000; Langridge and Struhl 2017), E-cadherin (Schwartz and DeSimone 2008) and Protocadherin-15 are involved in intercellular adhesions and transmission of mechanical stimuli. Protocadherin-15, for example, is involved in sensing sound vibrations across stereocilia tip links in the process of hearing (Kazmierczak et al. 2007). Mechanical forces are also known to be sensed at adhesions of cells with the ECM, as ECM stiffness dictates multiple cellular processes such as cell migration (Lo et al. 2000) and stem cell differentiation (Engler et al. 2006). For example, the ECM receptor CD44 is hypothesized to sense the stiffness of the ECM resulting in increased cell migration (Kim and Kumar 2014; Razinia et al. 2017). Additionally, the ECM receptor dystroglycan is thought to act as a shock absorber to protect the sarcolemma during muscle contraction (Barresi and Campbell 2006). Finally, even receptors that do not reside at canonical force sensing structures of cells have been implicated in mechanosensing. The RTK AXL which binds to a secreted ligand Gas6 has been shown to be a rigidity sensor (Yang et al. 2016) and facilitate a decrease in cellular stiffness in lung cancer (Iida et al. 2017). Thus, our studies showing that many receptors exhibit increased proteolysis in response to mechanical forces suggest that proteolysis may be a common mechanism used by cells to communicate mechanical stimuli. This assay could be used in the future to measure how varying the mechanical microenvironment affects receptor proteolysis.

### Limitations/caveats of assay

In the chimeric signaling assay, putative regions of proteolysis are evaluated in the context of artificial linkages at their N‐ and C-termini as well as potentially non-native stimuli and non-physiological presentation of proteases. In most cases, a small region of a receptor was excised and inserted into a larger receptor, resulting in non-native links to Notch’s ligand binding and transmembrane domains. One might expect the artificiality of the chimeras would result in a majority of chimeric receptors signaling either constitutively or not at all. However, several receptors exhibited “switch-like” behavior, underscoring the validity of SNAPS and the modular nature of cell-surface receptors. The use of the Notch transmembrane domain in the chimeric receptors also introduces some caveats as the domain, together with membrane proteins such as tetraspanins (Zimmerman et al. 2016), likely associates with the Notch membrane-tethered proteases ADAM10 and ADAM17. Though many of the chimeras studied have been reported to be cleaved by ADAM10 and ADAM17, some receptors may not typically reside in close proximity to these proteases and therefore not normally be cleaved. However, these proteases are upregulated in many diseases suggesting that the cleavage observed in this assay may be biologically relevant in certain cellular contexts. Finally, the chimeric Notch signaling assay provides a stimulus for exposing protease sites involving a 2-5 pN force normal to the cell surface. While many of the receptors studied here are also involved in cell-cell contacts likely involving similar mechanical forces, many interact with the ECM or have soluble ligands and perhaps may not normally be exposed to mechanical allostery. The main goal, however, was to provide a means to determine the presence of cryptic protease sites regardless of mechanical sensitivity. Harnessing this assay to study proteolytic regulation mechanisms is more specific than using, for instance, APMA to non-specifically activate metalloproteinases. (Ogata, Itoh, and Nagase 1995; Stetler-Stevenson et al. 1989)

## Conclusions

We have identified several putative proteolytic switches using SNAPS. These findings may drive development of conformation-specific modulatory antibodies as well as find use in synthetic biology applications that use cell to cell contact to drive transcriptional events. Our results provide a starting point for determining whether mechanisms of proteolytic regulation observed here are relevant to the biology of a given receptor. The convenient stimulus and response to proteolysis can be used to make additional chimeras to move closer to the native system and discover more about proteolytic regulation in the native receptor. For example, the luciferase response can be measured when systematically replacing chimeric domains with native transmembrane domains, ligand recognition domains, and intracellular tails. We also demonstrate that this assay provides a convenient platform for evaluating modulators of proteolysis.

## Acknowledgments

We would like to thank Steve Blacklow, Kassidy Thompkins, Maria Ramirez, and Robert Evans III for helpful comments on the manuscript and Klaus Lovendahl for cloning the TAM receptor chimeras and some of the RTK constructs. We thank the Aihara lab for use of their fluorescence plate reader. We would also like to thank Steve Blacklow and Jon Aster for the DLL4 stable cell lines and the Notch1-Gal4 construct. We would like to thank the Parker lab for the HER2 and HER4 cDNAs.

## Author contributions

W.R.G. and A.N.H. conceived and designed experiments. All authors analyzed data. A.N.H. and W.R.G. wrote the manuscript, and all authors edited. E.J.A. performed Herceptin and DECMA-1 antibody experiments. A.N.H., W.R.G., and E.J.A. prepared figures.

## Data and materials availability

All data needed to evaluate the conclusions of this study are available in the paper or the Supplementary Materials.

## Materials and Methods

### Reagents

Recombinant DLL4, MMP-2, and MMP-9 were purchased from R&D Systems. Batimastat (BB-94) was purchased from Sigma Aldrich. Compound E (GSI) was purchased from Fisher Scientific (Catalog # AAJ65131EXD). DECMA-1 antibody was purchased from Sigma-Aldrich (U3254). U2OS cells were purchased from ATCC. MS5 and MS5-DLL4 cells were a kind gift from Dr. Stephen Blacklow. 4-aminophenylmercuric acetate (APMA) was purchased from Sigma-Aldrich. Herceptin was purchased from MedChemExpress (HY-P9907). β-dystroglycan antibody was purchased from Leica Biosystems (B-DG-CE)

### Cloning

An Nhe1 site was added in Notch between amino acids 1735 and 1736 near the transmembrane region in a previously described Notch1-Gal4 construct (Andrawes et al. 2013) containing an N-terminal Flag tag, an AvrII site between the last EGF-like repeat and NRR, and a Bsu36i restriction site C-terminal to Notch transmembrane domain. All of the constructs were cloned using In-Fusion (Clontech).

CD44 was cloned using CD44S pWZL-Blast from Addgene (Item ID 19126). APP was cloned using pEGFP-n1-APP from Addgene (Item ID 69924). Dystroglycan was cloned from cDNA from Origene (Cat#: SC117393). mTFP was cloned from TS module from Addgene (Item ID 26021). AXL, MerTK, and Tyro3 were originally ordered as *E.coli* optimized gBlocks (IDT) for different constructs and then cloned into the Notch chimera using primers with In-Fusion ends. HER2 and HER4 DNA was a kind gift from Dr. Laurie Parker, from the ORF kinase library (Addgene). The remaining constructs were ordered as mammalian codon optimized gBlocks from IDT with In-Fusion ends.

### Cell culture

All cell lines were cultured in DMEM (Corning) supplemented with 10% FBS (Gibco) and 0.5% penicillin/streptomycin (Gibco). Cells were incubated at 37 °C in 5% CO_2_.

### Notch signaling assay

The Notch signaling assay was performed as described (Gordon et al. 2015). For co-culture assays, 0.1, 1, 2, or 10 ng chimera constructs were reverse transfected with reporter plasmids (50 ng Gal4 reporter plasmid and 1 ng PRL-TK reporter plasmid) in triplicate into U2OS cells in a 96-well plate. 24 hours post-transfection, MS5 cells or MS5 cells stably expressing DLL4 were plated on top of the U2OS cells with DMSO or drug (40 μM BB-94 or 1 μM GSI). 48 hours post-transfection, cells were lysed in passive lysis buffer (Promega). Lysate was added to a white 96 well half volume plate, and Dual-Luciferase Reporter Assay (Promega) was performed according to manufacturer’s recommendation and read out on a Molecular Devices LMaxII^384^ plate reader.

In assays using recombinant MMP-2 and MMP-9, activated MMP-2 or MMP-9 was diluted to 0.46 μg/mL in D10 media and added 36 hours post-transfection. Media was swapped out 38 hours post-transfection. Cells were lysed in passive lysis buffer 50 hours post-transfection and read out as previously described. For signaling assays using antibody, DECMA-1, Herceptin, or Sheep IgG control was added during the co-culture step 24 hours post-transfection.

### Cell surface ELISA

100 ng of Notch chimera constructs were transfected into U2OS cells in a sterile opaque tissue culture-treated 96 well plate (Corning 353296) in triplicate. 24 hours post-transfection, cells were washed once with PBS and fixed using 4% PFA (Thermo Fisher 28906) for 20 minutes, then washed three times with PBS. Cells were blocked in TBS+5% milk for 1 hour. Then, Flag primary antibody (Sigma-Aldrich F1804) was added 1:250 in TBS+5% milk for 2 hours. Cells were washed 3 times for 5 minutes each with TBS+5% milk. The cells were then incubated 1:10000 with an HRP secondary antibody for 1 hour before being washed 5 times for 5 minutes each with TBS. Chemiluminescent substrate was added for 1 minute before reading out on a luminescence plate reader.

### Western blot

48 hours post-transfection of dystroglycan constructs, Cos7 cells were lysed with RIPA buffer containing a protease inhibitor cocktail. Lysates were run on a 4-20% SDS-PAGE gel with 2 mM sodium thioglycolate in the running buffer. The protein was then transferred to a nitrocellulose membrane using a Genie Blotter (Idea Scientific) and blocked with 5% milk in TBS. β-dystroglycan antibody was diluted 1:1000 in TBS with 5% bovine serum albumin (BSA) added. A goat-anti mouse HRP conjugated antibody (Invitrogen) was used as a secondary antibody. Western blots were imaged using chemiluminescent buffer (Perkin Elmer Western Lightning Plus ECL) and the Amersham 600UV (GE) with staff support at the University of Minnesota-University Imaging Center.

## Supplementary Information

**Supplementary Figure 1-.**
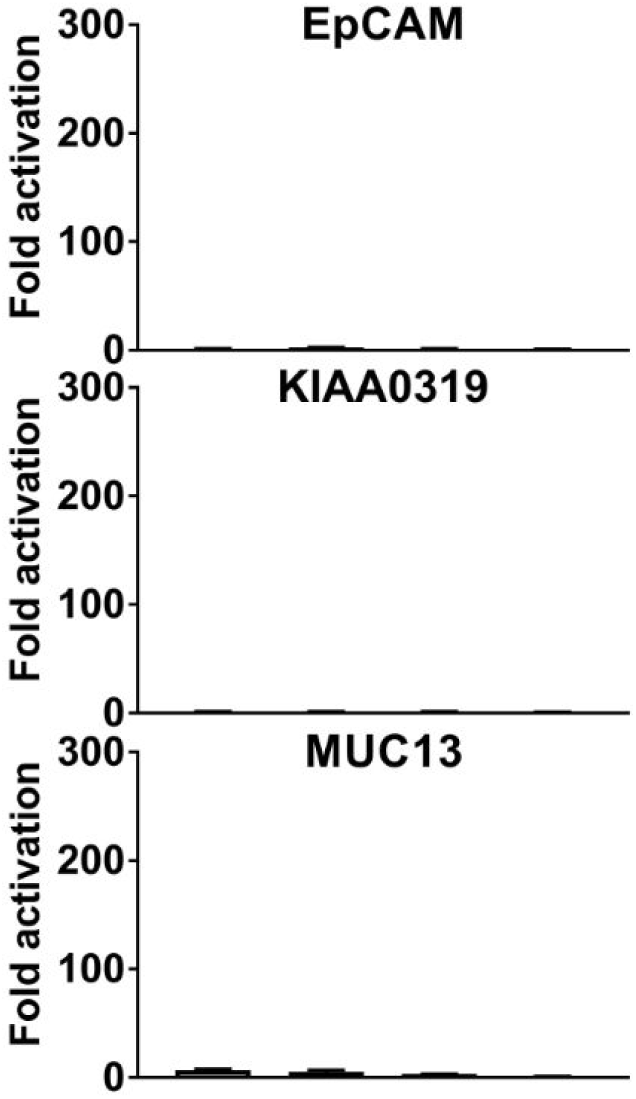
SEA domain chimeras without signaling activity. Luciferase reporter gene activity profile of Notch chimera constructs co-cultured with MS5 cells or MS5 cells stably transfected with DLL4, including treatment with BB-94 metalloprotease and GSI gamma secretase inhibitors as noted.

**Supplementary figure 2:**
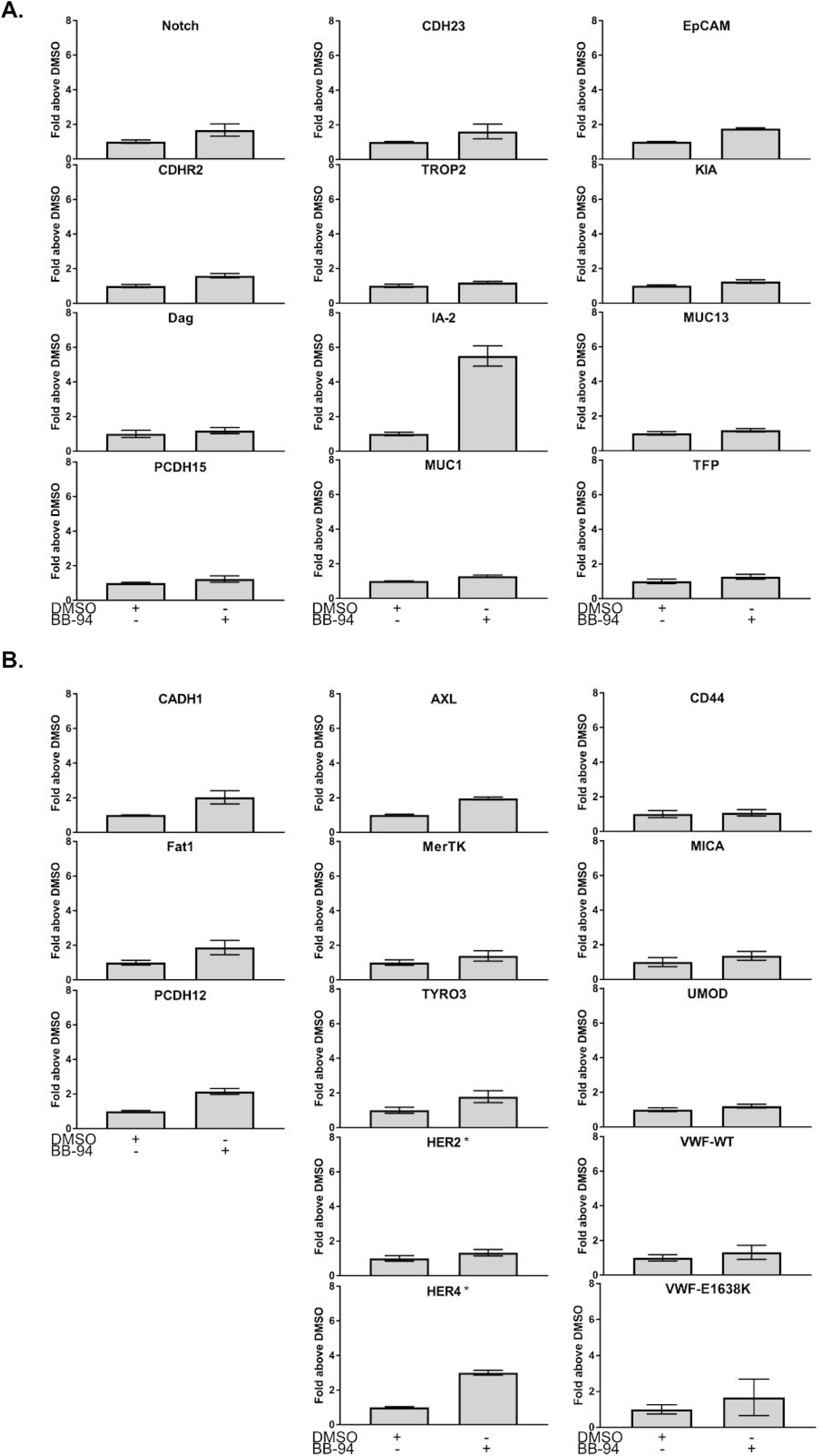
ELISA in the presence of BB-94. Cell surface ELISA performed with DMSO (negative control) or BB-94 (pan-metalloproteinase inhibitor). Data is normalized to the signal of DMSO condition. Error bars represent the SEM of triplicate measurements. A) SEA domains. B) Non-SEA domains. Asterisks denote cell surface ELISA was performed on a different date.

**Supplementary Figure 3:**
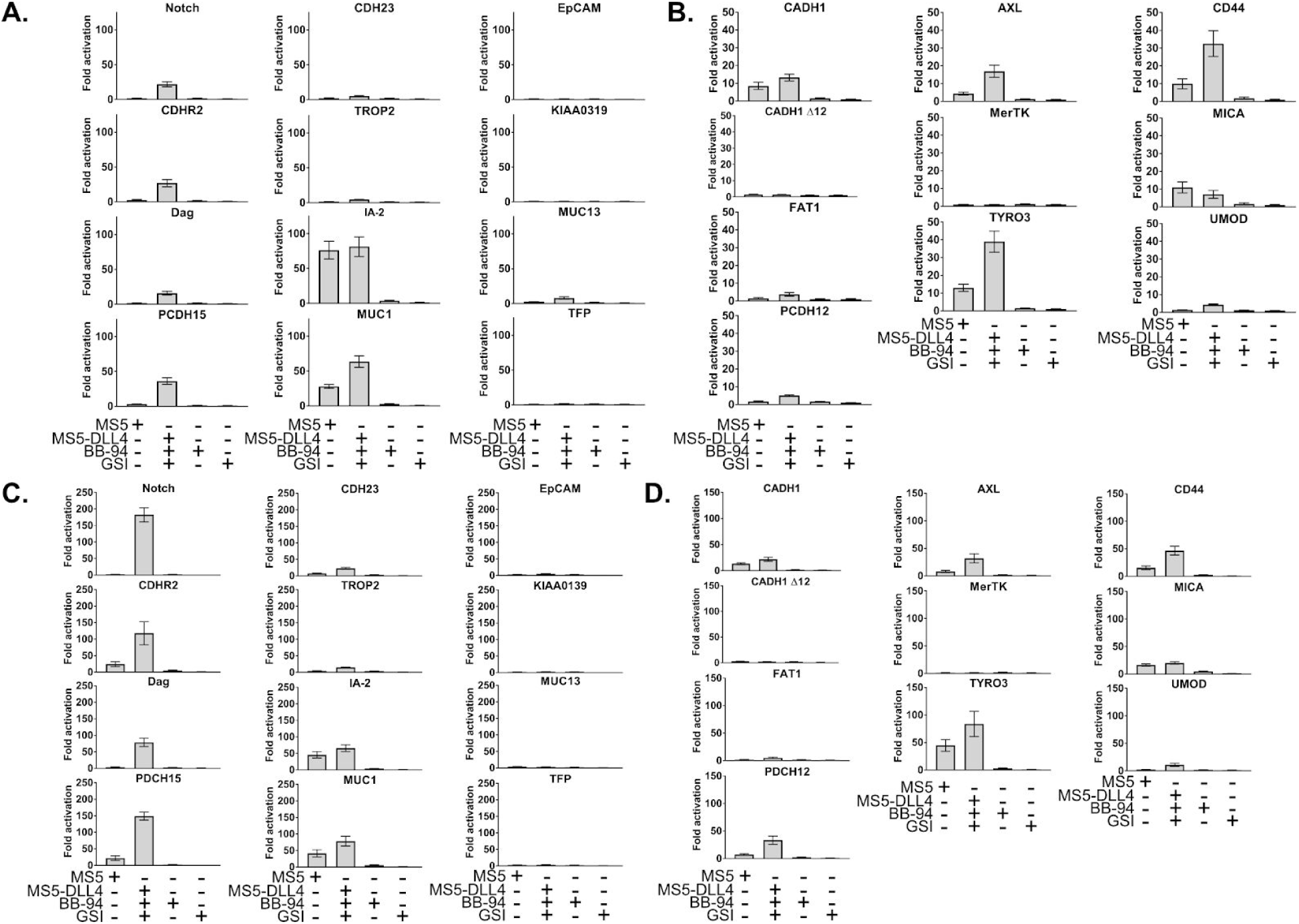
Titration of DNA used in co-culture assay. Luciferase reporter gene activity profile of Notch in comparison to the Notch chimera constructs with SEA/SEA-like domains (A and C) or diverse receptors (B and D) co-cultured with MS5 cells or MS5 cells stably transfected with DLL4. A) and B) 0.1 ng construct DNA per well. C) and D) 10 ng construct DNA per well. Error bars represent the SEM of triplicate measurements. BB-94=pan-metalloproteinase inhibitor GSI=γ-secretase inhibitor.

**Supplementary figure 4:**
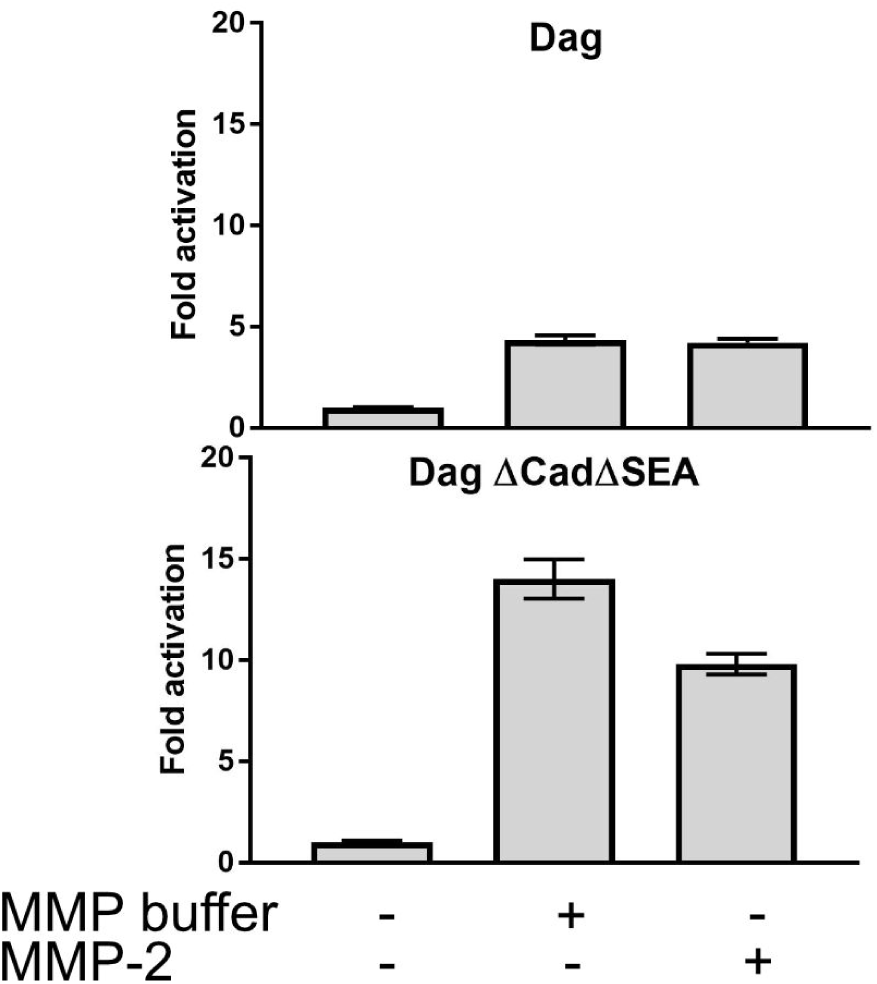
Comparison of MMP with MMP buffer. Notch chimera signaling assay using either activated MMP-2 or MMP buffer containing APMA. Error bars represent the SEM of triplicate measurements.

**Supplementary Figure 5:**
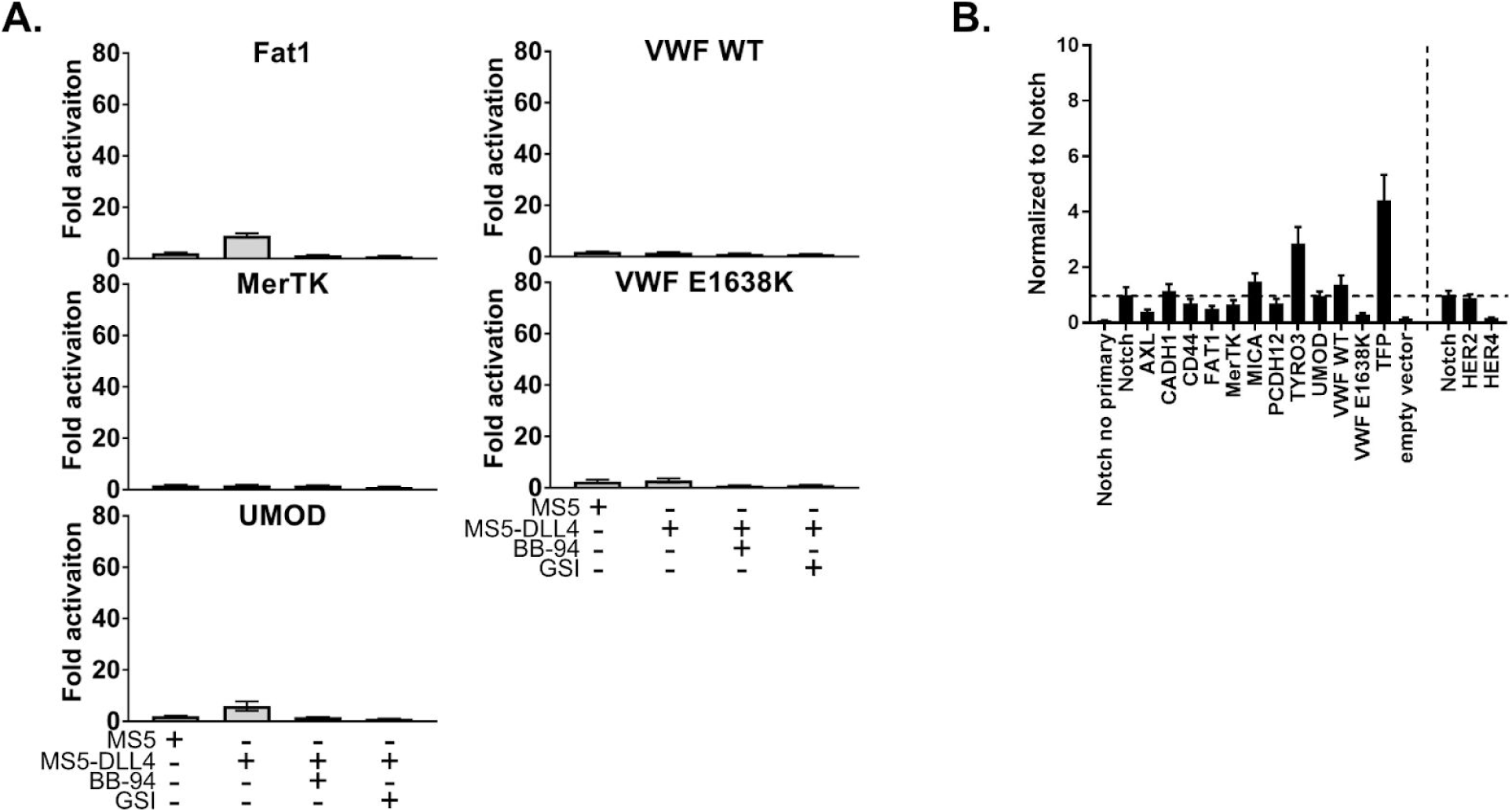
Additional data from diverse receptors. A) Luciferase reporter gene activity profile of diverse receptor chimeras co-cultured with MS5 cells or MS5 cells stably transfected with DLL4. B) Cell surface ELISA data from diverse receptor chimeras normalized to Notch signal. Horizontal dotted line denotes Notch signal, and the vertical dotted line separates cell surface ELISAs that were performed on different dates.

**Supplementary figure 6:**
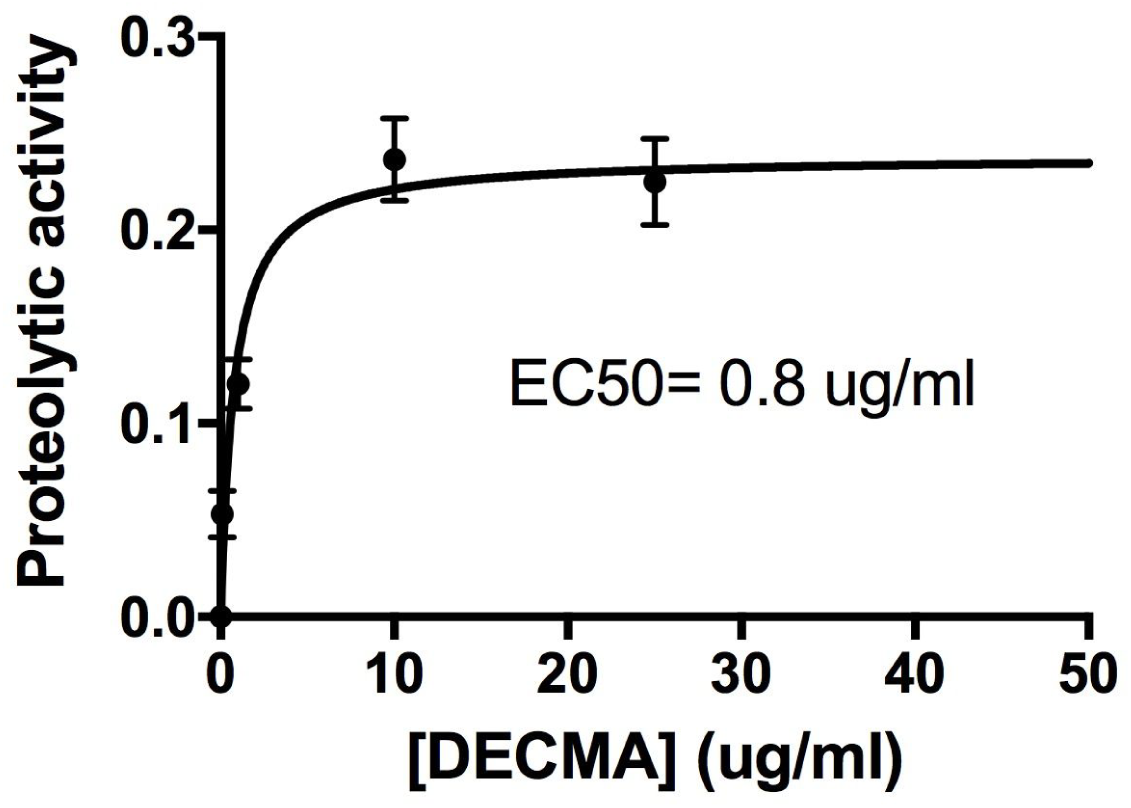
EC50 calculation for DECMA-1. The dose-dependent proteolytic activity enhancement of DECMA-1 was calculated by first subtracting the IgG control luciferase signal from the DECMA-1 signal at each concentration. The data was fit to a single exponential using Prism, including outlier detection.

**Supplementary Table 1:**
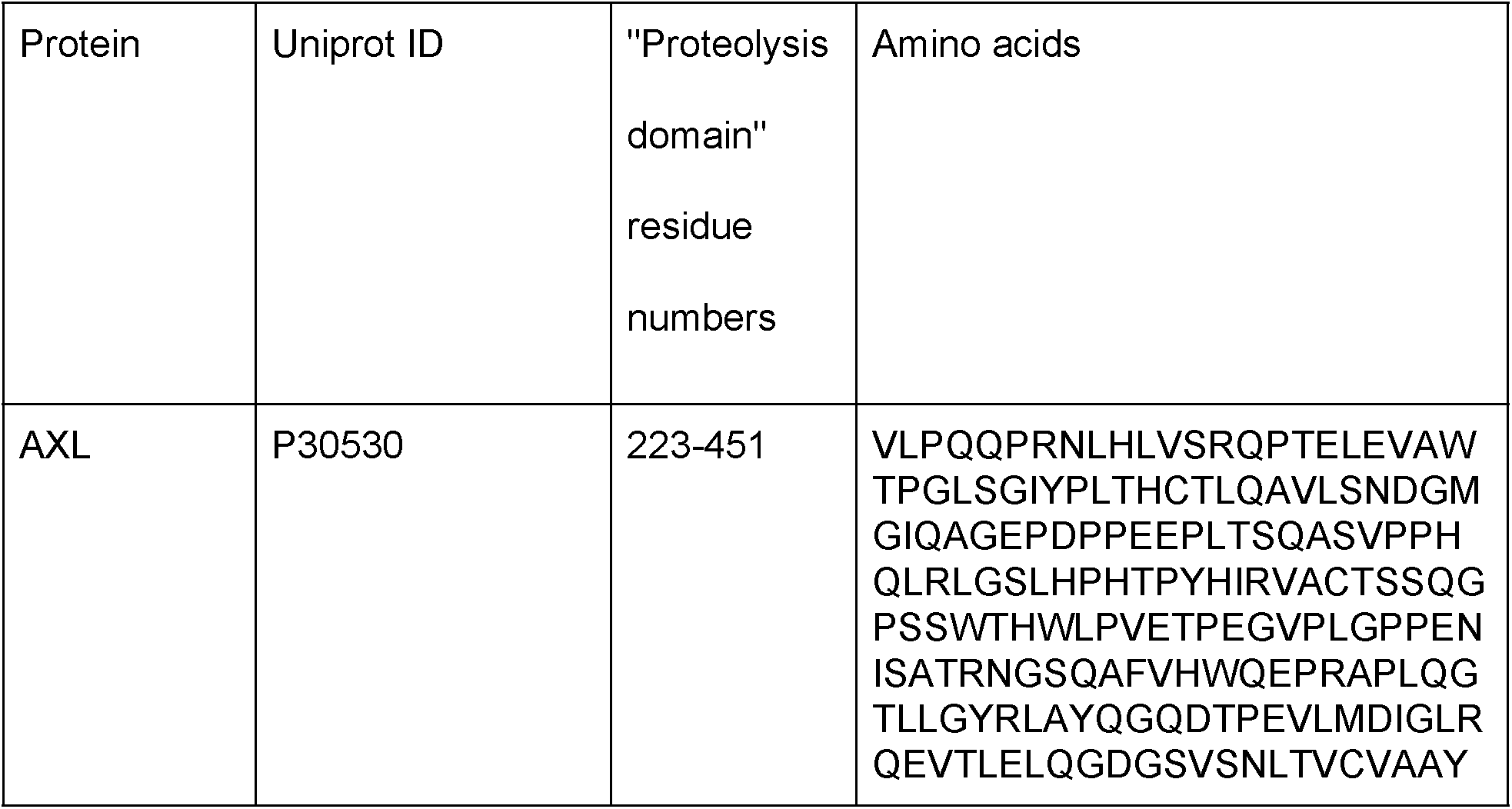
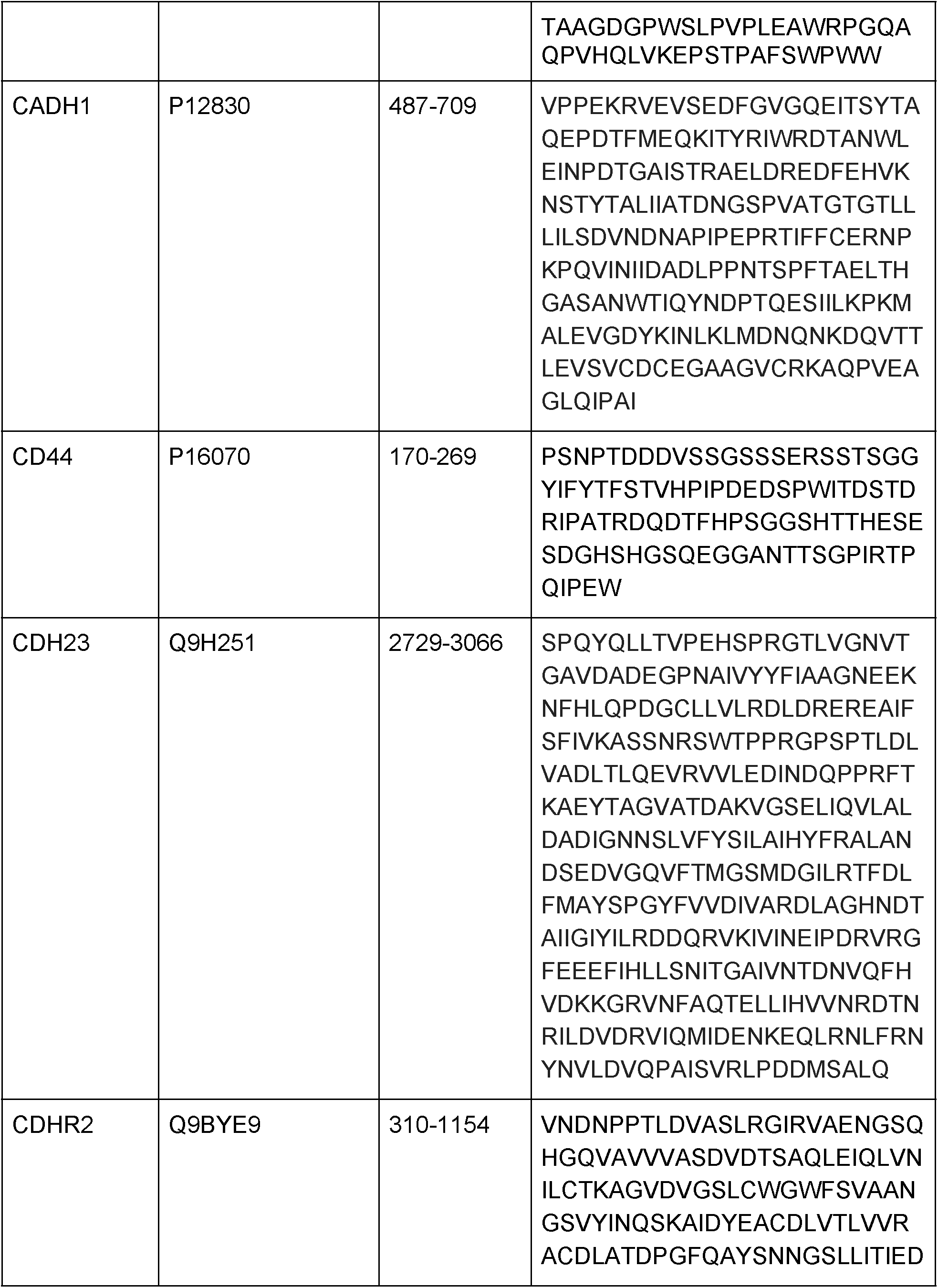
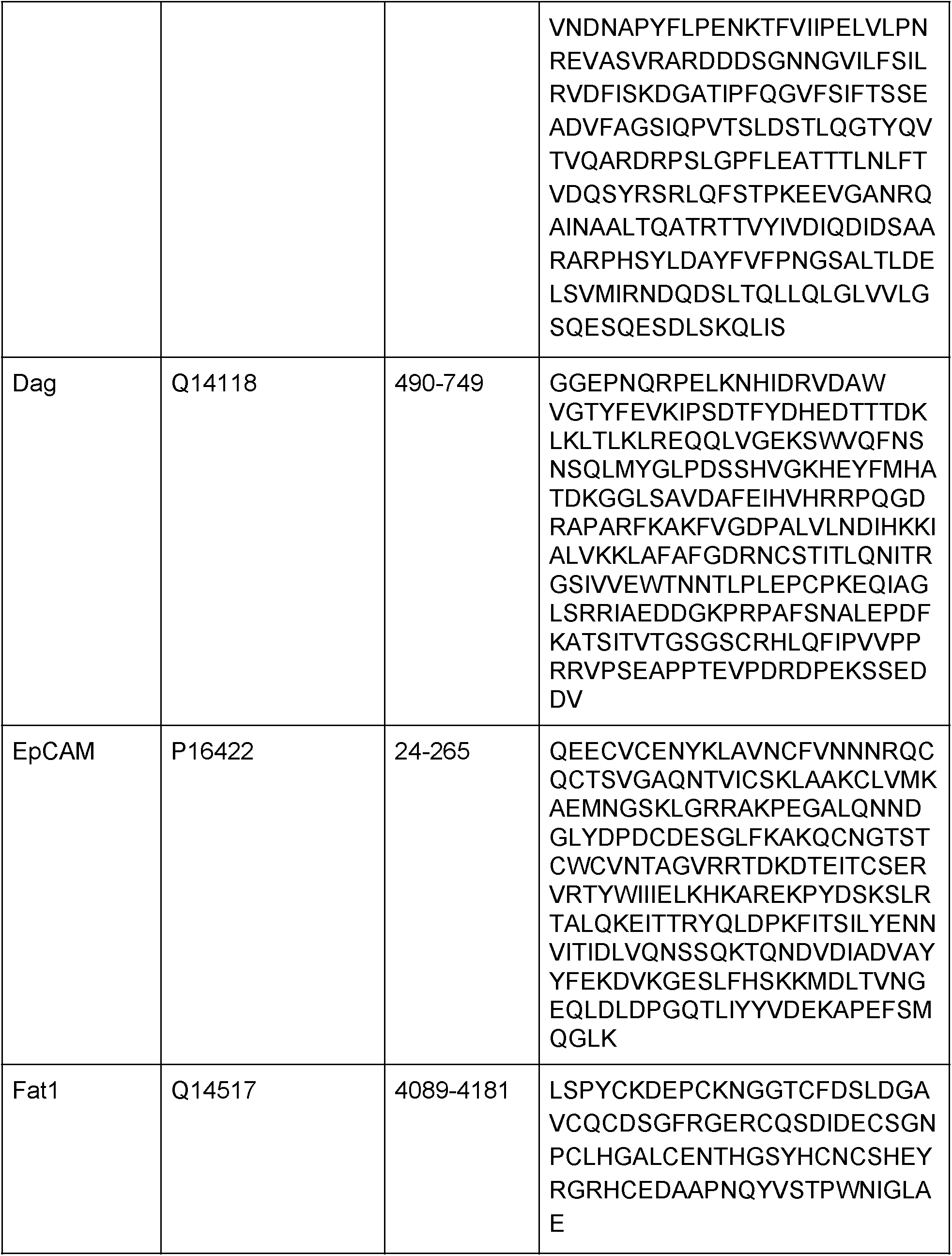
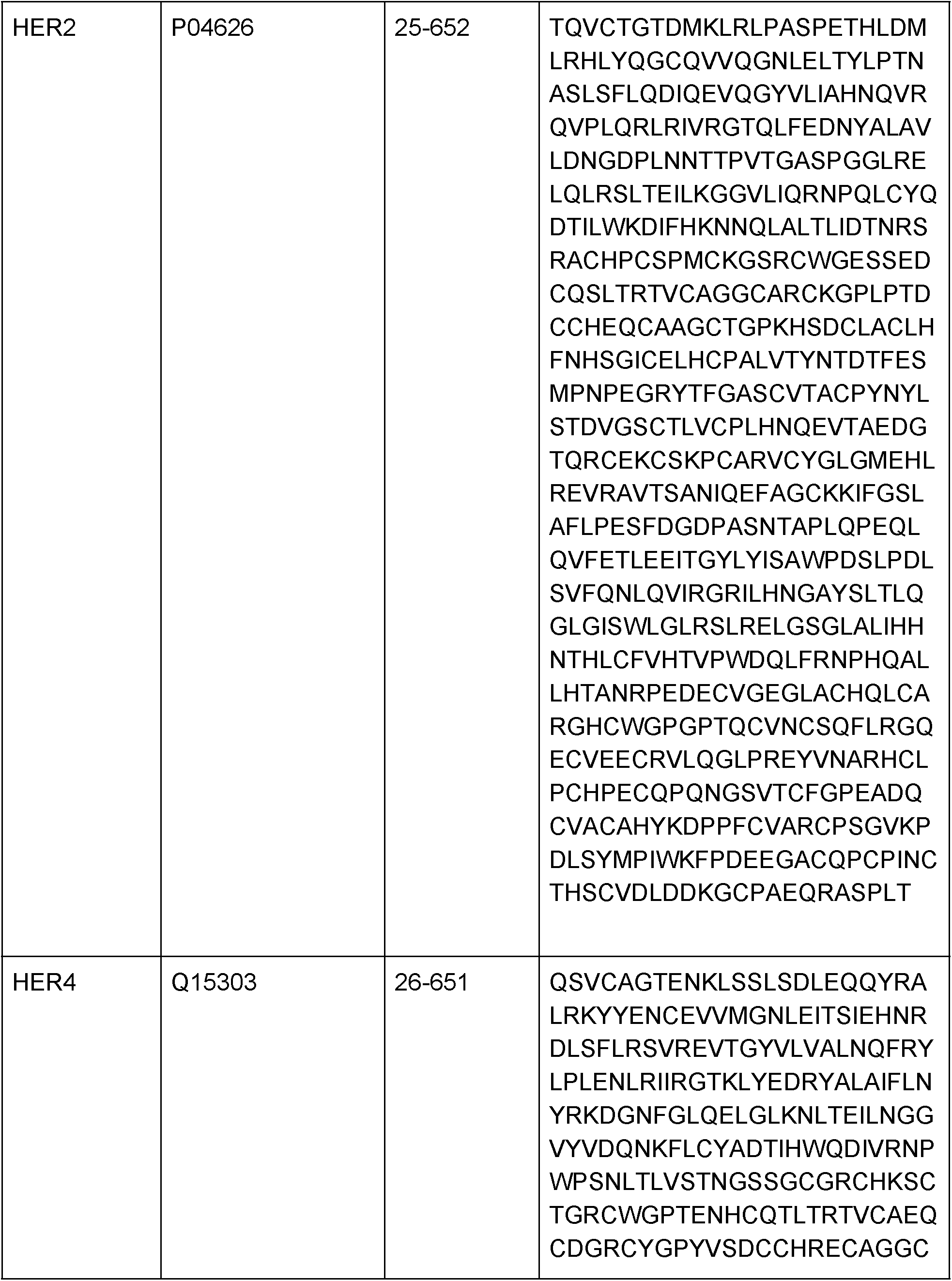
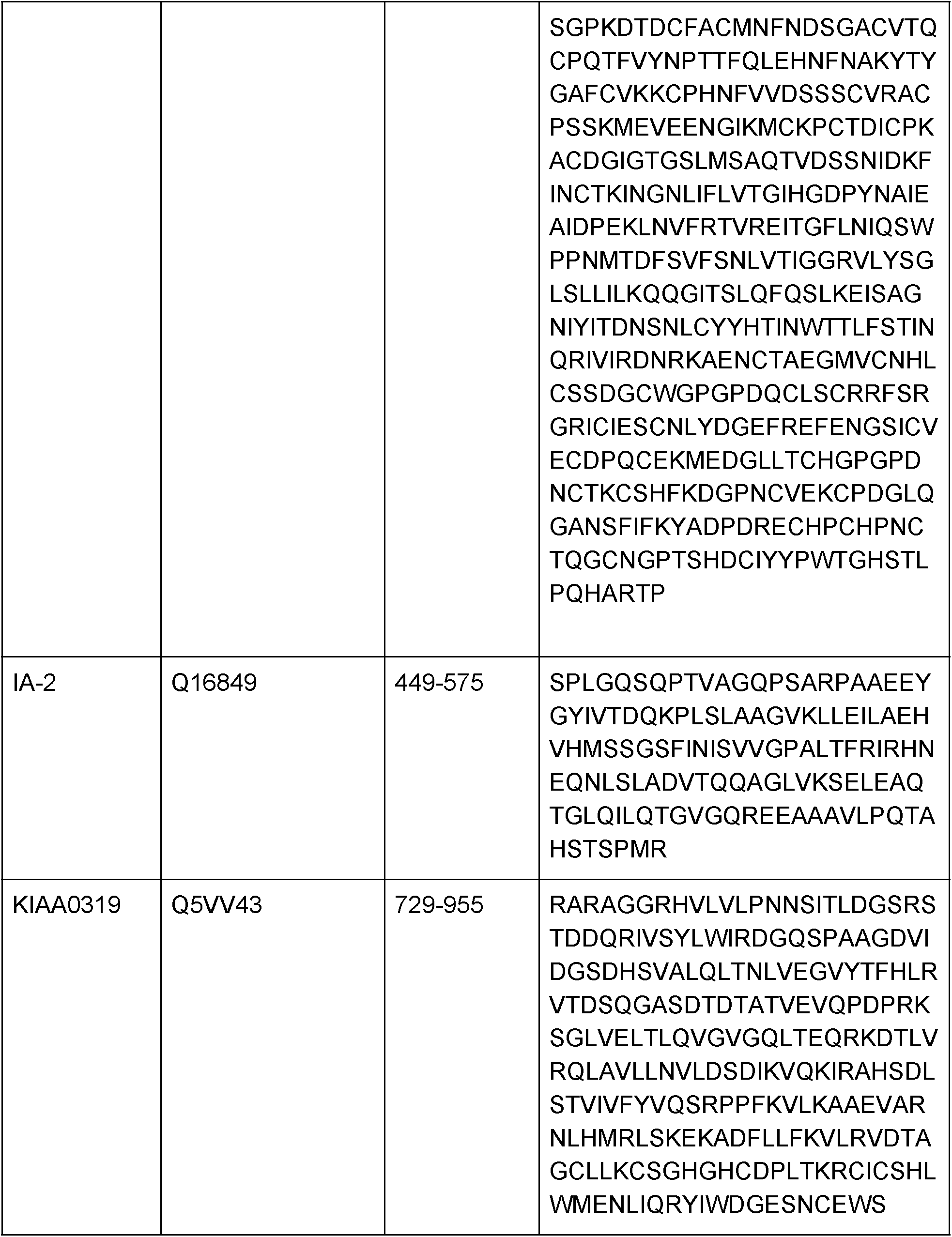
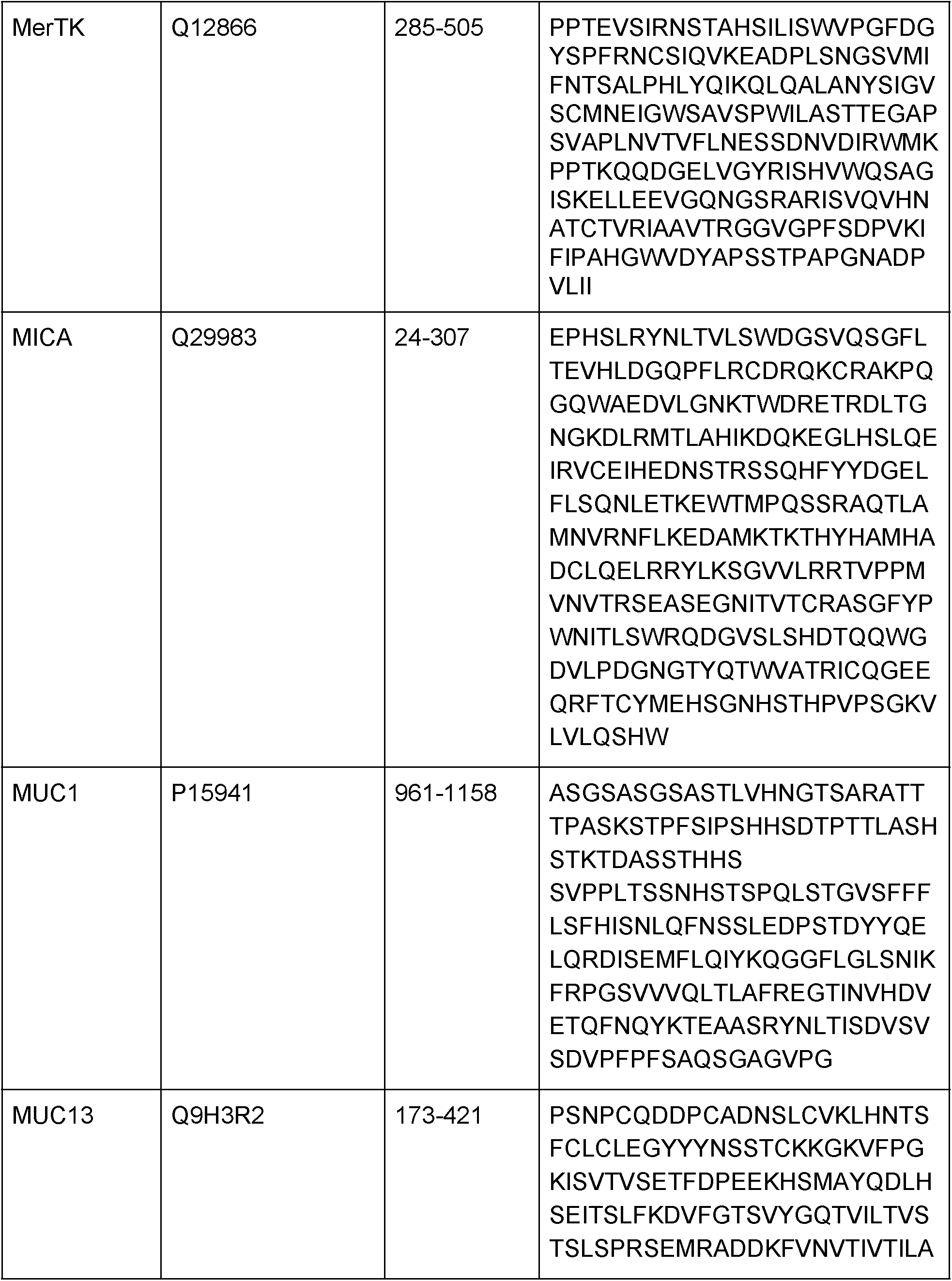
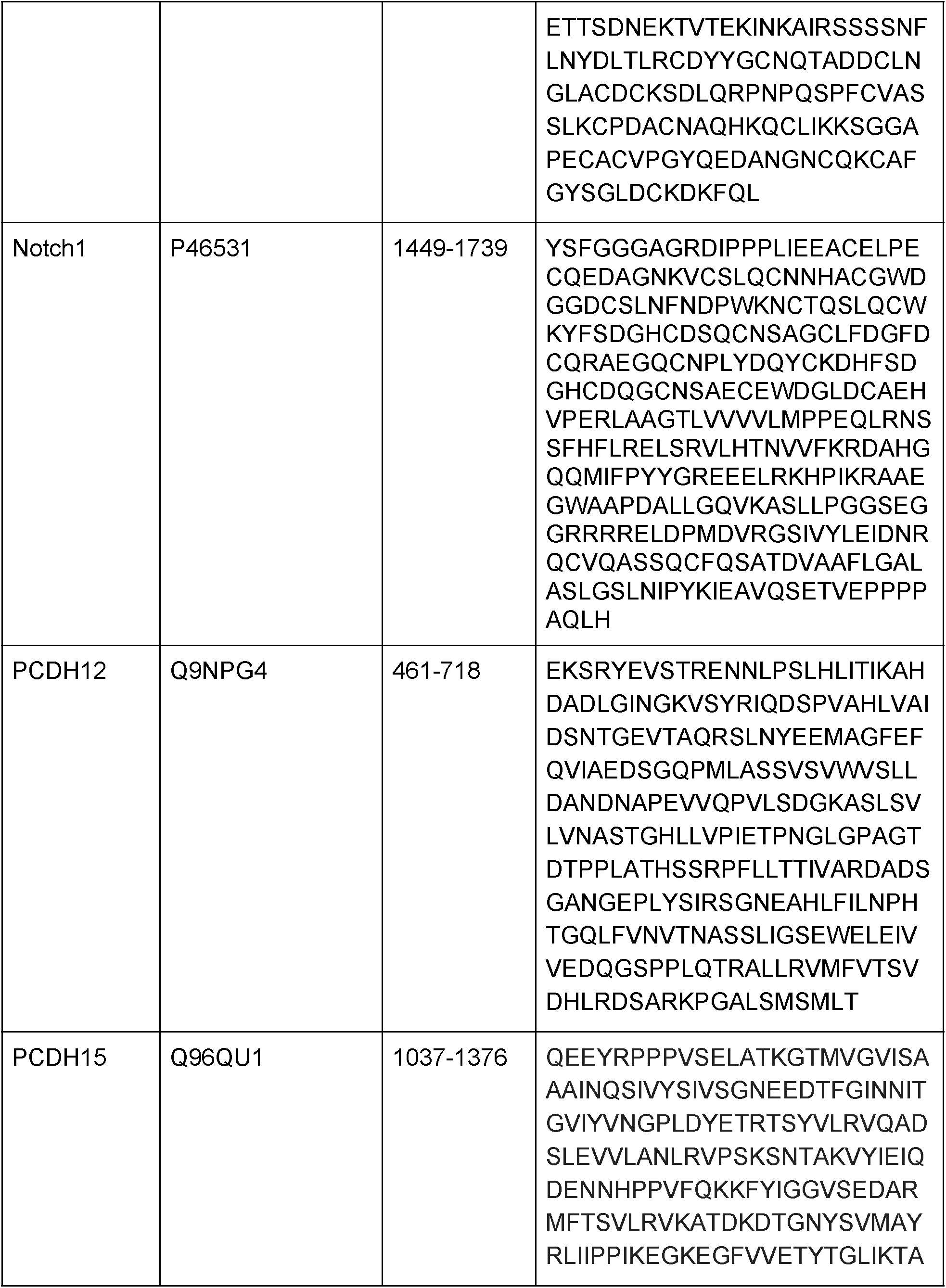
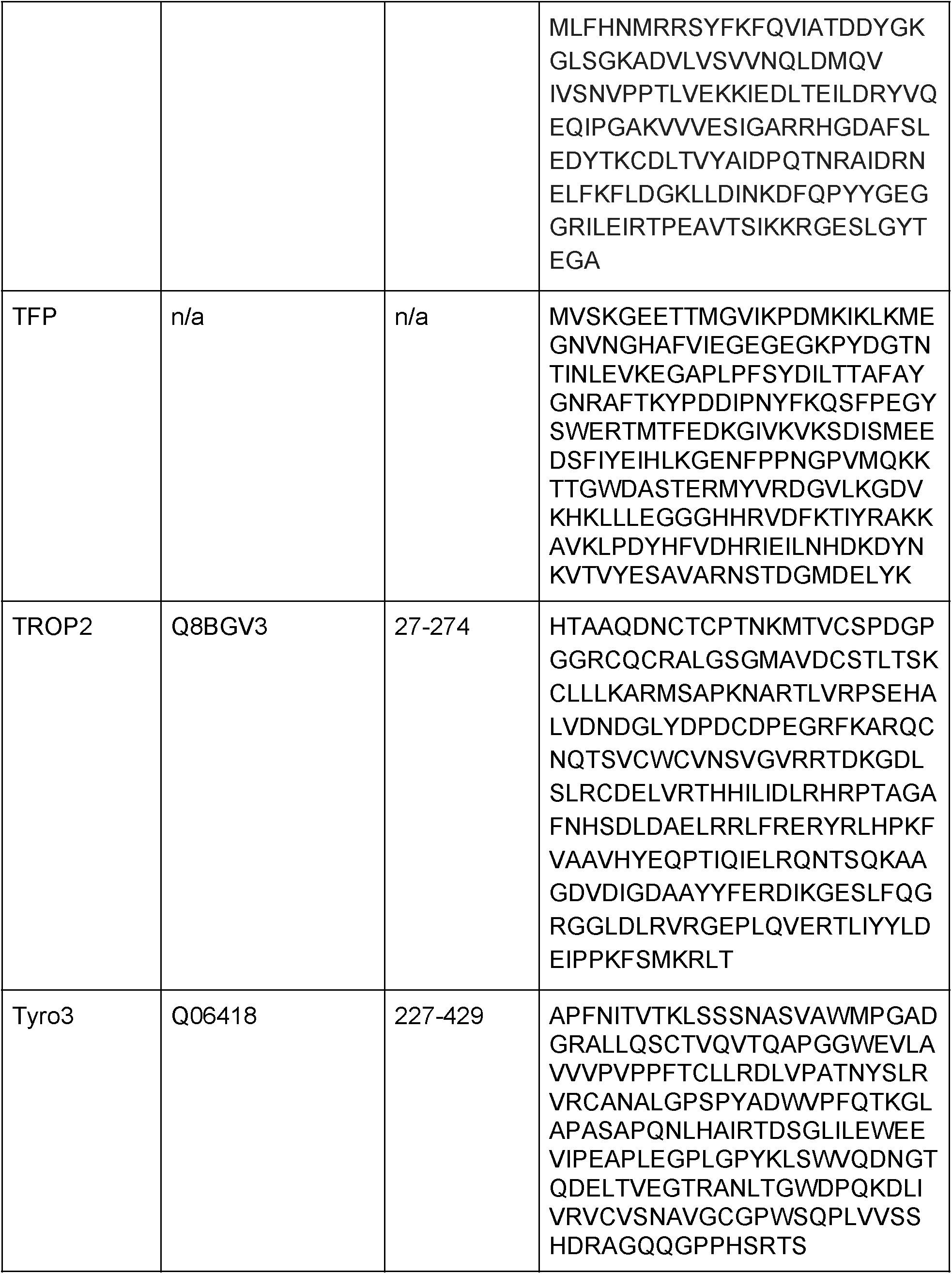
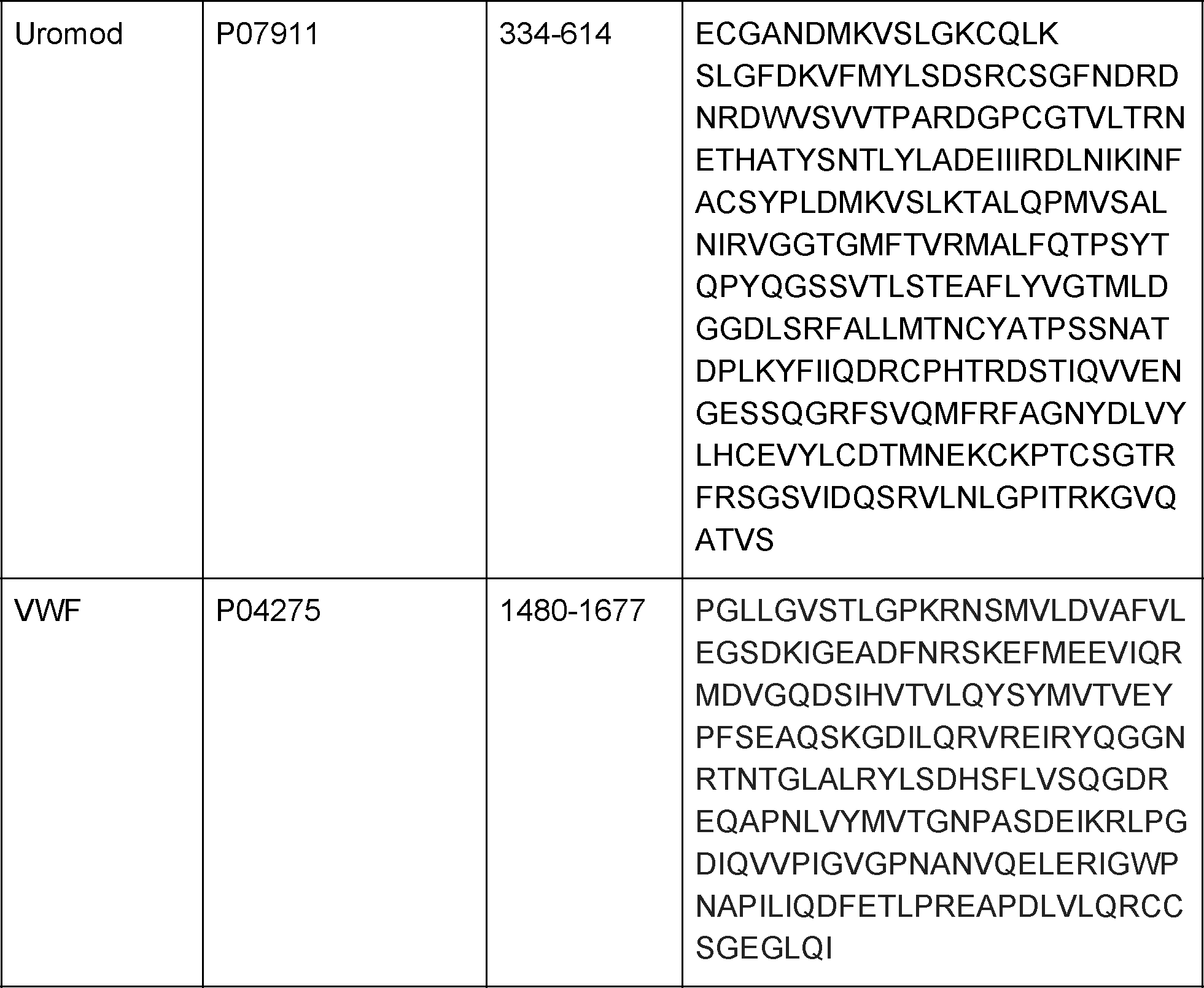
Sequences for all proteolysis domains used.

